# Container-based bioinformatics with Pachyderm

**DOI:** 10.1101/299032

**Authors:** Jon Ander Novella, Payam Emami Khoonsari, Stephanie Herman, Daniel Whitenack, Marco Capuccini, Joachim Burman, Kim Kultima, Ola Spjuth

## Abstract

**Motivation:** Computational biologists face many challenges related to data size, and they need to manage complicated analyses often including multiple stages and multiple tools, all of which must be deployed to modern infrastructures. To address these challenges and maintain reproducibility of results, researchers need (i) a reliable way to run processing stages in any computational environment, (ii) a well-defined way to orchestrate those processing stages, and (iii) a data management layer that tracks data as it moves through the processing pipeline.

**Results:** Pachyderm is an open-source workflow system and data management framework that fulfills these needs by creating a data pipelining and data versioning layer on top of projects from the container ecosystem, having Kubernetes as the backbone for container orchestration. We adapted Pachyderm and demonstrated its attractive properties in bioinformatics. A Helm Chart was created so that researchers can use Pachyderm in multiple scenarios. The Pachyderm File System was extended to support block storage. A wrapper for initiating Pachyderm on cloud-agnostic virtual infrastructures was created. The benefits of Pachyderm are illustrated via a large metabolomics workflow, demonstrating that Pachyderm enables efficient and sustainable data science workflows while maintaining reproducibility and scalability.

**Availability:** Pachyderm is available from https://github.com/pachyderm/pachyderm. The Pachyderm Helm Chart is available from https://github.com/kubernetes/charts/tree/master/stable/pachyderm. Pachyderm is available out-of-the-box from the PhenoMeNal VRE (https://github.com/phnmnl/KubeNow-plugin) and general Kubernetes environments instantiated via KubeNow. The code of the workflow used for the analysis is available on GitHub (https://github.com/pharmbio/LC-MS-Pachyderm).

## Introduction

The relevance of big data in biomedicine is evident. Technological advances in fields such as massively parallel sequencing [1], mass spectrometry [2] and high-throughput screening [3] are examples of how biology has shifted towards a data intensive field [4]. The rapid increase in the number of data points and the size of the observations in those fields pose many difficulties, but this is definitely not the only obstacle. Apart from the need to process large amounts of data, computational biologists must manage analyses that include multiple stages and tools, while simultaneously maintaining reproducibility of results.

Undoubtedly, a major concern to the scientific method is that results should be fully reproducible by other researchers. Using multiple distinct processing tools makes it hard for scientists to replicate results [5]. One appealing option is to use workflow systems such as Snakemake [6], Galaxy [7] or Nextflow [8]. These systems help to coordinate complex dependencies between tools, aiding researchers with their analytical duties. Nevertheless, scientific workflows constructed in one environment are not likely to easily run on other environments straightaway [9].

Previously, virtual machines (VMs) were introduced as a feasible approach to achieving system-agnostic deployments. However, this approach has numerous disadvantages; given the large size and poor ability to reuse software components inside of VMs. The microservices-based architecture offers a compelling alternative with the possibility to divide complex applications into a collection of smaller, more focused services that communicate with technology-agnostic protocols [10]. Software containerisation provides an ideal solution with frameworks such as Docker (https://www.docker.com) to enable microservices based architectures [11]. Containers isolate an application’s view of the underlying operating environment including all the required packages and libraries. In contrast to virtual machines, containers can be launched in a relatively short time period, by avoiding the installation of redundant dependencies on host machines, and the need to boot the guest operating systems. Their lightweight nature makes them very flexible and particularly well suited for computations in cloud environments because far more computing units (containers) can be deployed on demand. The microservice architecture is gaining importance within science, as it provides better reproducibility and standardisation of computer-based analyses [11]. Thanks to containerisation, scientists can package pipelines in an isolated and self-contained manner, to be distributed and run across a wide variety of computing platforms. Examples of projects in which microservices are a cornerstone include the PhenoMeNal project [12] and the EXTraS project [13].

An important component for microservices is the need for a framework to orchestrate them on multiple compute nodes. Kubernetes is an open source project coordinated by Google for automating deployment, scaling, and management of containerised applications [14]. This technology is able to coordinate a cluster of interconnected computers to work as a single unit, by automating the distribution and scheduling of containers across the cluster. Pachyderm (http://www.pachyderm.io/) is a workflow and data management tool built on top of Kubernetes. This framework is capable of running a piece of analysis in parallel over a set of containers in a cluster, promising good scalability. It also addresses interoperability and reproducibility through the containerisation of software tools and a fully versioned file system. There is a wide range of existing workflow tools like Bpipe [15] and Reflow (https://github.com/grailbio/reflow) that allows users to manage data pipelines, but most are tied to specific languages (e.g., Python) and/or are not designed with native support for containers. There are also existing tools like the Dat Project (https://datproject.org/) or Git Large File Storage (https://git-lfs.github.com/) that provide data management or versioning, but these tools are not natively integrated with any kind of pipelining or processing tools. iRODS [16] can in effect be used to implement both a file access layer and a form of pipelining solution through its powerful rule language. Though, its rule language is not developed specifically for the needs of reproducible data processing, which makes this solution more complex than using a dedicated pipelining framework. Pachyderm is unique in that it manages pipelines and the associated data in a unified and reproducible manner, with pipelines treated as a first class citizen. Certainly, it is one of the few workflow tools, together with Argo (https://applatix.com/open-source/argo/), built as a layer on top of Kubernetes, such that it is completely portable and language/data agnostic. Despite being relatively new, Pachyderm has been used in a number of different settings, including bioinformatics use cases such as germline variant calling and joint genotyping (https://github.com/pachyderm/pachyderm/tree/master/doc/examples/gatk), as well as in sensor analytics (https://www.microsoft.com/developerblog/2017/04/12/reproducible-data-science-analysis-experimental-data-pachyderm-azure).

The main focus of this project is to demonstrate the efficacy of Pachyderm in the context of bioinformatics data processing. In particular, we will present how Pachyderm enables scalable, reproducible, and portable bioinformatics workflows, using a metabolomics workflow as an example. Moreover, we will show how Pachyderm can be deployed on any Kubernetes-based infrastructure by means of a Helm Chart we created, and how it is possible to extend the Pachyderm File System to work with traditional block storage.

## System and methods

### Kubernetes

Kubernetes is a tool that efficiently orchestrates containerised applications in a cluster of computers, without tying containers specifically to individual machines. Kubernetes provides different abstractions for defining workloads such as pods, jobs, services or replication controllers. Amongst these abstractions, pods are the essential units of computing that can be created and managed in Kubernetes. A pod can be defined as a group of containers and volumes that are bundled and scheduled together because they share resources like a file system or a network address. In order to simplify complex operations with Kubernetes resources, Helm (https://helm.sh/) is available as a package manager for versioning, releases, deployments, deletions and updates of containerised applications.

### Pachyderm workflow tool

Pachyderm is a platform for managing data pipelines and the associated input/output data in a manner that results in all data pipelines being reproducible and scalable. To enable its functionality, Pachyderm leverages projects from the container ecosystem including Docker, Kubernetes, and etcd (https://coreos.com/etcd). These projects allow Pachyderm to be language and framework agnostic because the units of data processing are defined by software containers.

To use a software tool within Pachyderm, researchers simply need to write the code associated with a pipeline processing stage in a preferred programming language and make sure that it can read and write files from and to a local file system. They then need to, via a JSON pipeline specification, declaratively supply Pachyderm with a name for the processing stage, a Docker image in which the code will run, a command to execute in the running container, and one or more data input(s). Subsequently, Pachyderm will ensure that the corresponding containers are run on an underlying Kubernetes cluster, and it will inject the input data that needs to be processed into the running containers. In cases where a processing stage is specified to run in parallel, Pachyderm will share the input data across running containers and collect the corresponding outputs.

The main element of Pachyderm is pachd, which is the Pachyderm daemon, or server, that manages all of the pipelining and data versioning features of Pachyderm. This daemon runs in one or more pods in Kubernetes and communicates with users and/or other components of the system via GRPC (https://grpc.io/). Figure 1a gives an overview of the different components of pachd, including a pipeline system, a file system and a block store component.

**Figure 1:**
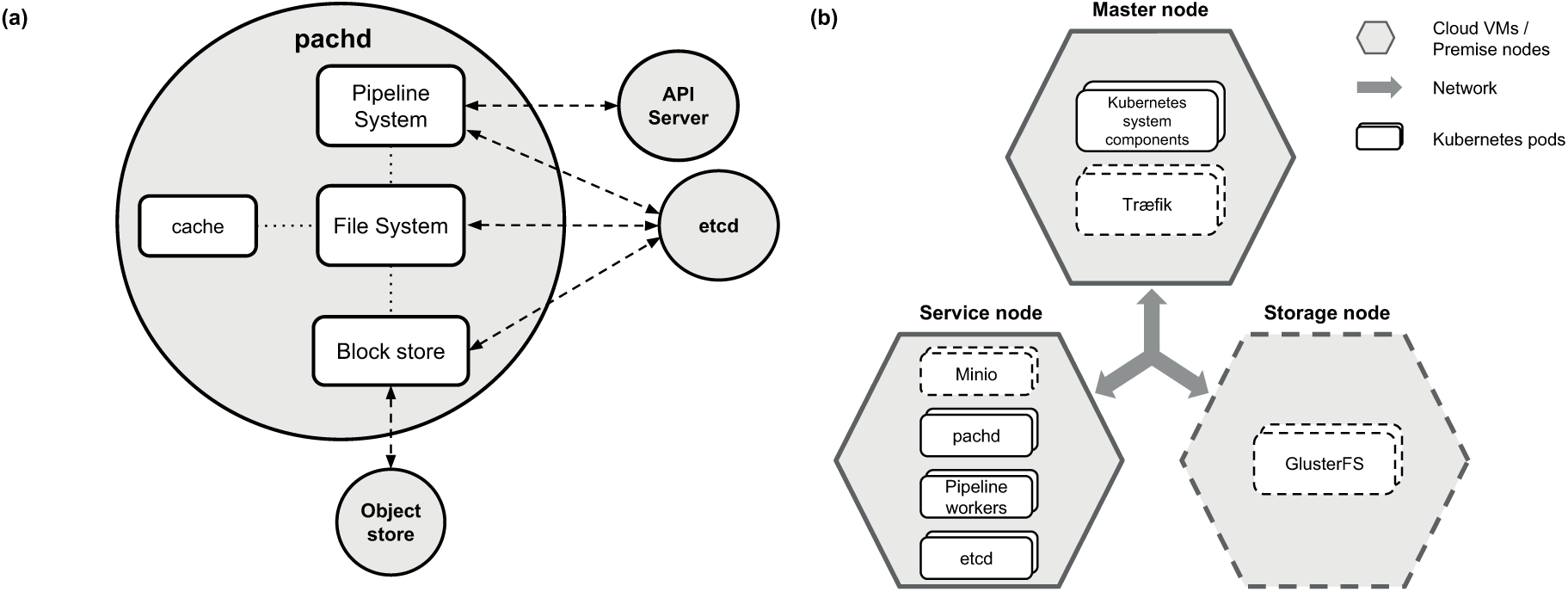
(a) The Pachyderm daemon. pachd is the Pachyderm daemon managing the pipelining and data versioning features of Pachyderm. The main components of pachd are (i) a file system component, (ii) a block store component, and (iii) a pipelining component. The file system component handles all requests related to putting data into and getting data out of Pachyderm Data Repositories (PDRs). To this end, the file system component cooperates with the block store component to content address new data, put new objects in the backing object store, pull objects out of the backing object store, etc. The pipelining system component creates and manages all of the pipeline workers, which execute to process data in Pachyderm pipelines. The pipelining system component cooperates with the file system component to make sure that the correct subsets/versions of data (versioned in PDRs) are provided to the correct pipeline workers, such that data is processed in the sequence and manner specified by users. To coordinate and track all of these actions, pachd stores and queries metadata in etcd, a distributed key/value store that is also deployed in a pod on Kubernetes, and it communicates with the Kubernetes API Server and the backing object store service. Further, Pachyderm optimises uploads/downloads of data via an internal caching system. **(b) A typical infrastructure and services setup with Pachyderm.** A standard Kubernetes cluster contains two major entities represented in two different polygonal figures. Cloud virtual machines/premise nodes are depicted as hexagons, whereas Kubernetes pods are displayed as rounded rectangles. Optional nodes/pods are depicted with dashed borders. The master node coordinates the rest of the nodes, runs the Kubernetes API and can use a reverse proxy such as Træfik (https://traefik.io/). In the service nodes, all Pachyderm related pods are scheduled: the Pachyderm daemon, Pachyderm pipeline workers and etcd. Also, Minio services can be deployed in service nodes, responsible for upload/download of data to/from the backing storage. The storage dedicated node (optional) is in charge of providing application containers with a shared file system (e.g. GlusterFS), using block storage volumes.

### Pachyderm File System (PFS)

The Pachyderm File System component of pachd utilises a copy-on-write paradigm which is based on Git-like semantics. It manages (i) the versioned data repositories, (ii) aids in shimming data into containers for processing, and (iii) stores newly committed input or output data into the backing object store. When one or more files and/or directories are committed into a Pachyderm data repository, the PFS content addresses each input file to create a hash. The files are stored in the backing object store via this hash, and the file system represents this hash in etcd, where the repositories, jobs, and provenance of the data are tracked.

All input/output data managed by Pachyderm is organised into versioned collections of data called Pachyderm Data Repositories (PDRs). This concept of versioned data repositories combined with the pipelining system gives rise to Pachyderm’s unique concept of data provenance. Data provenance, also referred to as data lineage, refers to the metadata associated with the origin, evolution and movement of the data over time [17]. In the PFS, any particular state of data can be identified by commits. Pachyderm can give users all the data repository names, commit IDs, and versioned pipeline specifications corresponding to specific states of data. Thus, any run of any pipeline producing any result is completely reproducible and explainable, at least in terms of the corresponding data transformations and intermediate states of data.

When the pipelining system indicates that data needs to be processed, the file system component, along with a binary that is injected into the user container, retrieves the relevant data from the object store and shims it into the user container under a local directory “/pfs/<name of the input repository>”. The file system also collects any results written to “/pfs/out” by the user container and automatically versions these into a corresponding output data repository.

Regarding storage options, the PFS can be backed in any of the generic object stores provided by Google Cloud Platform (https://cloud.google.com/), Amazon Web Services (https://aws.amazon.com/), Microsoft Azure (https://azure.microsoft.com) and Minio (https://minio.io/).

### Pachyderm Pipeline System (PPS)

In Pachyderm, data processing is performed by pipeline workers, which can be thought of generally as a user’s Docker containers running their code. However, to be more specific, the workers are Kubernetes pods. The pods include Docker containers based on the user-specified Docker images along with Pachyderm components, where the Pachyderm components support the triggering and data management logic discussed below.

The pods are created when the corresponding pipeline is created and are, by default, left running in the cluster waiting for new data to be available for processing. As new "commits" of data are made on the repositories that are specified as input to pipelines, Pachyderm stores that data and triggers the corresponding processing stage performed by the pipeline workers. In many cases, only the newly added/modified data is supplied to the workers, such that the processing occurs incrementally. This means that instead of processing all results every time, Pachyderm is able to reuse previous results and compute only what is necessary, making it resource efficient. In Figure 2, an example of how the PFS and PPS utilise the two most characteristic features of Pachyderm can be observed: data provenance and incremental processing.

**Figure 2:**
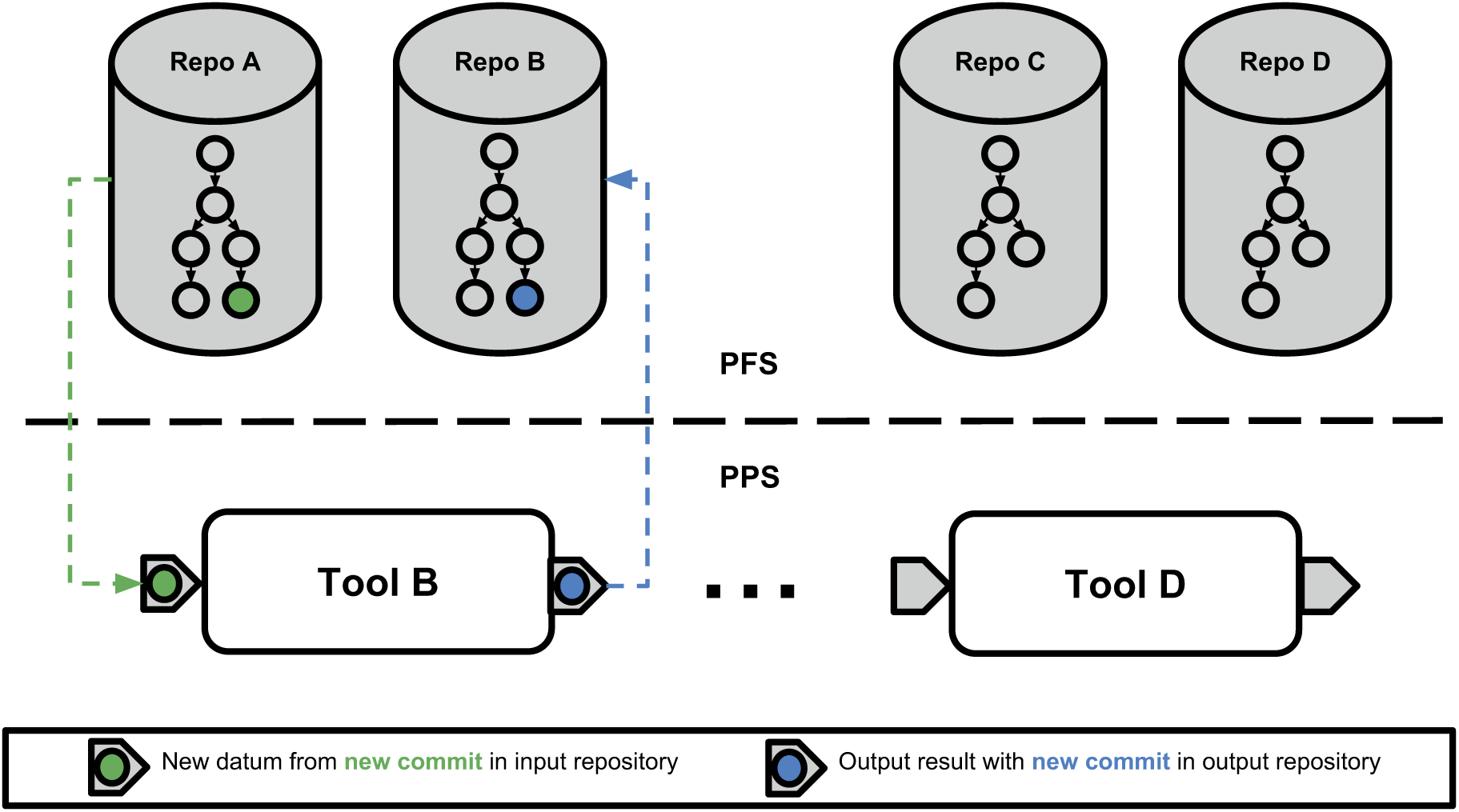
Data provenance and incremental processing in Pachyderm. The upper part of the figure shows the Pachyderm Data Repositories (PDRs) present in the Pachyderm File System (PFS) after creating a bioinformatics workflow. These repositories contain a tree-like structure in which each node represents a separate commit. In the lower part of the figure, the different pipeline stages of the workflow are displayed in the Pachyderm Pipeline System (PPS), together with their corresponding inputs and outputs. When a new data commit (green color) is added to the input data Repo A, a new pipeline stage is triggered for Tool B in the PPS, leading in turn to a new commit (blue color) in Tool B’s output repository. Thereby, the provenance of the blue commit made on Repo B would be: (i) the green commit from Repo A and (ii) Tool B’s pipeline specification. Note that the commit structure looks similar for the two data repositories because of the nature of a linear data pipeline. In the figure, the repos created by Tool B and Tool C do not have the corresponding commits as the data processing has not yet reached this level in the pipeline. As new commits are added into the PFS, the PPS triggers the corresponding pipeline stages with the new datums (minimal computing units) from the commit. This phenomenon can be referred to as incremental processing, as only new computing units are processed. These new datums are then computed, creating further commits on downstream repositories and providing a mechanism to track the provenance of the computations.

Once a pipeline worker completes the processing of new data commits, the pipeline worker gathers any data written by the containers to "/pfs/out" and versions that data in a PDR corresponding to that particular workflow stage. Thereafter, another stage can specify that output as its input. In this way, it is possible to declaratively define directed acyclic graphs (DAGs) of processing, which are driven by data repositories and the commits of data corresponding to those data repositories.

Pachyderm can perform parallel computations by partitioning the data into various subsets called "datums" which are the minimal data units of computation. The contents of these datums are user-defined glob patterns. For example, a glob pattern of "/" would tell Pachyderm that it always has to process all of the input repository together as a single datum, whereas a glob pattern of "/*" would tell Pachyderm that it could process any files or directories in the root of the repository in isolation (as separate datums).

PPS then works together with PFS to spread out the datums across the pipeline workers. The number of pipeline workers that can complete this work in parallel is user-defined and can be constant or a coefficient of the cluster size. Each of the pipeline workers downloads and processes one datum at a time in isolation, and then pachd gathers all of the results corresponding to each datum, combines them together, and versions this combination as the output of the pipeline. The scheduling of pipeline workers is determined by the load on the various nodes and the resource requests (i.e., for memory, CPUs, GPUs, etc.) that are made for each pipeline. This allows Pachyderm to optimally utilise the underlying resources.

In summary, the operations involved in running a multi-stage Pachyderm workflow are:

1. Create input PDRs for the workflow
2. Create a first workflow stage that processes those input PDRs
3. Create one or more other workflow stage(s) that processes the output of the first stage and/or other downstream stage(s)
4. Commit data into the input PDRs

Once the above sequence of steps is completed, Pachyderm will automatically trigger the necessary "jobs" to process the input data, which will include a job to process the data in the input PDRs along with any dependent downstream jobs. Each of these jobs will run to either success or failure. In the case of a success, the job will make an output commit of results into the PDR corresponding to the pipeline. In the case of a failure, no output commit of data will be made from the pipeline.

## Results

In general, bioinformatics researchers have access to cloud environments that use different interfaces to their resources, varied architectures and implementation technologies. This makes it important for solutions to be cloud agnostic, so they can be compliant with diverse infrastructures (e.g. private Openstack [18] environments). Further, it is crucial to provide an easy to follow procedure to instantiate virtual infrastructures of multiple nodes, and install and configure frameworks without much expertise. One attractive way of solving this is to use Virtual Research Environments (VREs) to aid the deployment and configuration of complete virtual infrastructures with the associated software for data analysis. Here, we present a convenient solution so that scientists can incorporate Pachyderm into their infrastructure, regardless of their cloud-provider and storage backend. Furthermore, we demonstrate by means of a metabolomics case study how Pachyderm can enable scalable and sustainable workflows.

### Extending the PFS for block storage

The Pachyderm File System is not POSIX-compliant, as it relies on an object storage backend. The PFS can be backed by S3 compliant object stores, Blob Storage in Azure or GCS in Google Cloud. Unfortunately, these object storage options are not always available on all infrastructures. As an example, the infrastructure used for the case study in the manuscript only supports block storage, which is managed by a shared file system (GlusterFS). In order to overcome these limitations, we extended the Pachyderm File System to work with block storage. This was achieved by enabling Minio to act as an object store interface with our storage backend. Thanks to Minio, it is possible to add a highly available, load-balanced S3 object-store compatibility to the storage tier of block storage based infrastructures.

### Pachyderm Helm Chart

In order to deploy Pachyderm in multiple settings, we developed a Helm Chart that makes the workflow tool entirely cloud-agnostic, and most interestingly, makes it easier to deploy it backed in multiple storage options. Thanks to this chart, users can easily install Pachyderm on Kubernetes-based infrastructures from any cloud provider, such as Openstack. Besides, it provides scientists with a flexible and straightforward mechanism to configure various settings of Pachyderm, such as the resource requests and the storage backend used by the Pachyderm File System.

Currently, the Helm Chart supports five general deployment scenarios, which include: (i) local deployment on Minikube, (ii) on-premise deployment, (iii) Google Cloud, (iv) Amazon Web Services and (v) Microsoft Azure. As an example, an Openstack user should opt for an on-premise installation, which is compatible with any Cloud infrastructure. This type of deployment makes Pachyderm completely cloud-agnostic, as the only requirement is that it necessitates a S3 endpoint as storage backend, like Minio. The chart has been pushed to and is now maintained on the official Kubernetes Charts repository (https://github.com/kubernetes/charts/tree/master/stable/pachyderm). In addition, Pachyderm is available out-of-the-box from the PhenoMeNal VRE (https://github.com/phnmnl/KubeNow-plugin) and general Kubernetes environments instantiated via KubeNow [19]. The latter makes it straightforward to launch a complete virtual infrastructure with Kubernetes and Pachyderm installed on the major cloud providers.

### A metabolomics case study

The study of metabolomics concerns the comprehensive profiling of low-weight molecules, known as metabolites, comprising the metabolomes of e.g. biological specimens. As metabolites are the intermediate and end products in all biological pathways, changes caused by various pathophysiological processes will immediately impact the metabolome, thereby making it an attractive target for biomarker discovery [20]. Liquid chromatography coupled to mass spectrometry (LC-MS) has gained momentum in the field as it provides large amount of information about the specimens in a relatively short time. In fact, a modern mass spectrometer is able to produce 35K spectra per hour [21], emphasising the need to adopt automated and scalable approaches in order to effectively process the generated data.

For this case study, a dataset (containing 138 LC-MS runs) was used that included 37 cerebrospinal fluid (CSF) samples that were measured in duplicates. Twenty-seven of the samples originated from patients diagnosed with multiple sclerosis (MS) of which 13 depicted a relapse-remitting phenotype (RRMS) and the remaining 14 a progressively degenerative phenotype (secondary progressive MS, SPMS). The dataset also contained measured CSF metabolomes of ten non-MS and non-inflammatory controls. The dataset is available in the MetaboLights database [22] (MetaboLights ID: MTBLS558, http://www.ebi.ac.uk/metabolights/MTBLS558).

The objective of creating a metabolomics workflow in Pachyderm was to demonstrate a real-world scenario in which patients’ data are processed in a scalable, interoperable and reproducible manner. We implemented a computational workflow to process LC-MS data, illustrated in Figure 3, and evaluated how well it can scale on a Kubernetes infrastructure. The workflow has been described thoroughly elsewhere by Khoonsari et al. [12]. Briefly, the open source mzML files were first centroided and calibrated using OpenMS [23]. To this end, the signals resulted by each metabolite were clustered to so called mass traces that were used for quantification. This clustering was performed using OpenMS (FeatureFinderMetabo) and XCMS (findPeaks) [24]. The mass traces were corrected for retention time drift and were matched across the samples using group and retcor functions in XCMS. The mass traces were filtered based on presence/absence in the blank samples as well as correlation to dilution series. The resulting mass traces were grouped and annotated with adduct information using CAMERA [25]. For identification, the *MS*^2^ data was read and mapped to adduct information to calculate neutral mass of the precursor ions. This information was then used in CSI:FingerID [26] to identify metabolites using database searching. Finally, multivarate statistical analysis was performed on the identified metabolites using partial least squares discriminant analysis (PLS-DA) [27].

**Figure 3:**
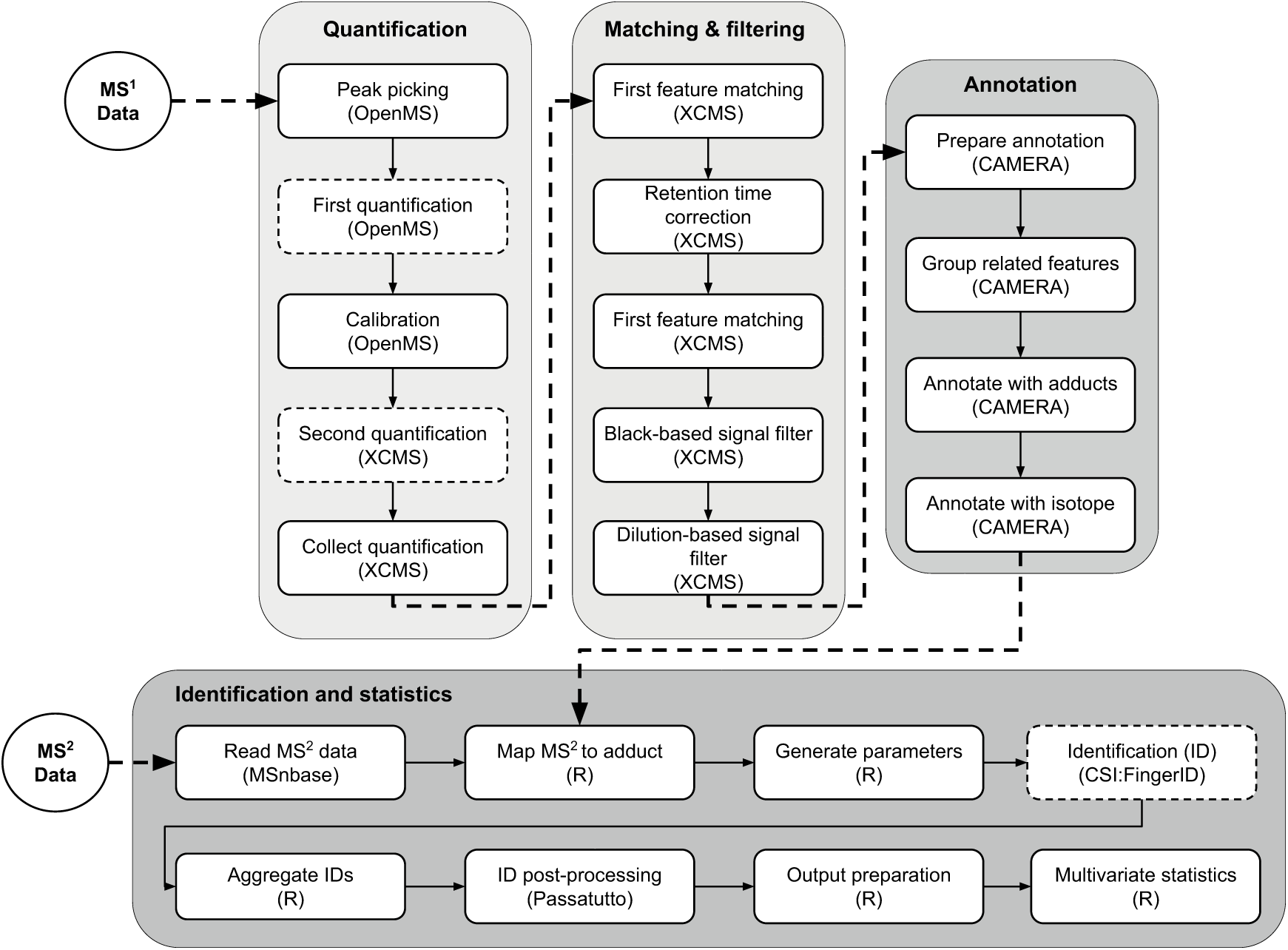
LC-MS workflow definition. The workflow consists of five main components including quantification, matching and filtering, annotation, identification and statistics. The raw *MS*^1^ data in open source format (e.g., mzML) is accepted as input. In the quantification component, the raw data is first centroided, calibrated and the signals from each metabolite are clustered into mass traces. In the matching and filtering component, the retention time drift is corrected and the mass traces are matched across the samples. The non-biologically relevant signals are filtered based on presence/absence in blank samples as well as correlation to dilution series. In the annotation component, the mass traces are annotated with adduct and isotope information. This information is used in the identification component to calculate the neutral mass of the precursor ions. The identification is then performed and the resulting scores are converted to posterior error probability values. The data is then limited to the mass traces annotated with an identification hit and subjected to multivariate data analysis. Note that the pipeline stages chosen for the performance benchmarks are illustrated with dashed borders.

### Infrastructure setup

We set up a Kubernetes cluster on the Amazon Web Services cloud using KubeNow. The cluster was composed of a master node, several service nodes and a number of storage nodes. All of the nodes had the same flavour (t2.2xlarge) with 8 vCPUs and 32GB of RAM. The number of storage nodes and service nodes varied, as we scaled the infrastructure to evaluate its performance with different numbers of work executors.

### Services setup

The Minio object store service (release 2018-01-02T23-07-00Z) was deployed in shared mode using the official Kubernetes Minio Helm Chart (https://github.com/kubernetes/charts/tree/master/stable/minio). The number of Minio replicas used equals to half the number of workers used for the analyses. Similarly, Pachyderm (version 1.7.0rc2) was deployed using our developed (and now the official) Pachyderm Helm Chart. An on-premise deployment mode was performed, using one replica, 1 CPU and 3GB of RAM requests for pachd and etcd, and 5GB of PFS cache. A more detailed description of a typical services and infrastructure setup is depicted in Figure 1b.

### Performance

There are several metrics for measuring parallel scaling performance. Two of these methods are speedup and scaling efficiency [28]. Both are important measurements in Cloud Computing, since resources are commonly pay-per-use. The speedup gives an estimate on how much faster computations are performed using a larger number of workers, whereas the scaling efficiency tells how efficient computations are when increasing the number of parallel processing elements. We studied the speedup and scaling efficiency of the three most CPU-intensive and parallelisable jobs in the LC-MS workflow (OpenMS’s FeatureFinderMetabo, XCMS’s findPeaks, and CSI:FingerID) calling attention to the number of processing units used during different runs. FeatureFinderMetabo and findPeaks were applied on each sample (n=138), while CSI:FingerID was applied on each *MS*^2^ spectrum extracted from the eight *MS*^2^ samples (n=5000).

The speedup (*S*(*N*)) can be specified as *T*_1_/*T_N_*, where *T*_1_ is the serial running time, and *T_N_* is the multiple worker parallel running time, being *N* the number of processing elements (e.g., number of PPS workers). In order to obtain *T*_1_, the median of the serial processing times of the runs of each tool was calculated. It is worth noting that measured serial processing times do not include upload/download times, whereas parallel processing times do. This gives a better representation of the time that tools take on a single-core environment with data locally available. Also, parallel processing times include the container instantiation time, but not the time needed to pull the required container image, as container images are usually previously pulled. The scaling efficiency can be defined as the ratio of speedup to the number of processing elements.

For each tool studied, we benchmarked its performance against the speedup and scaling efficiency using different numbers of workers and different cluster settings. Table 1 presents an overview of the cluster setup used for our performance analysis.

**Table 1:** Cluster setup used in the benchmarks

| Service nodes | Storage nodes | Minio replicas | Workers |
| --- | --- | --- | --- |
| 5 | 3 | 10 | 20 |
| 8 | 4 | 20 | 40 |
| 11 | 6 | 30 | 60 |
| 14 | 7 | 40 | 80 |

The results obtained after evaluating the speedup and scaling efficiency of the three different tools are presented in Figure 4, see also Supplementary Table 1. The first benchmark was executed using OpenMS’s FeatureFinderMetabo tool, which took ∼37.3 minutes to run when using 80 PPS workers. The serial running time of this tool summed up to ∼30.8 hours, resulting in a speedup of ∼50 and a scaling efficiency of 62%. Using the same cluster setup, we found a speedup of ∼55 and a scaling efficiency of 69% when evaluating XCMS’s findPeaks tool. In this case, the serial running time was ∼10.5 hours, while a parallel running time of ∼11.5 minutes was obtained. The last benchmark was carried out using CSI:FingerID annotation tool. It took ∼10.1 minutes to run it when using 80 PPS workers. The serial running time of this tool was ∼10.9 hours, leading to a speedup of ∼63 and a scaling efficiency of 79%. As shown in Figure 4, the scaling efficiency decreases with a larger number of workers.

**Figure 4:**
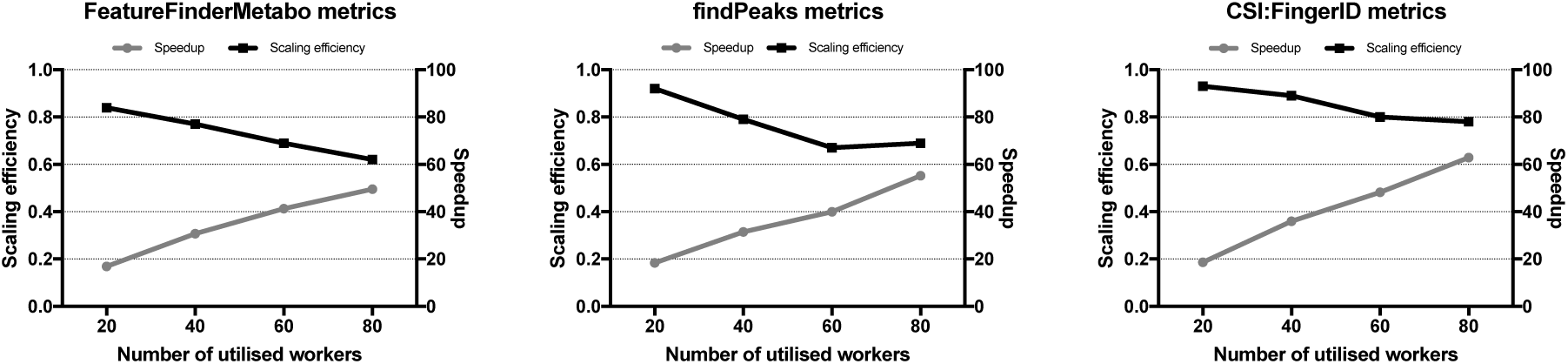
Performance metrics. Each of the figures displays the speedup (right axis, grey line) and scaling efficiency (left axis, black line) obtained when utilising various numbers of workers with three different tools of the metabolomics workflow.

## Availability

Pachyderm is open source and available on GitHub (https://github.com/pachyderm/pachyderm). We created a Helm Chart that is available on the official Kubernetes repository (https://github.com/kubernetes/charts/tree/master/stable/pachyderm). Pachyderm is available out-of-the-box from the PhenoMeNal VRE (https://github.com/phnmnl/KubeNow-plugin) and general Kubernetes environments instantiated via KubeNow. The code of the metabolomics workflow used for the analysis is available on GitHub (https://github.com/pharmbio/LC-MS-Pachyderm).

## Discussion

The goal of this study was to demonstrate Pachyderm as a potential bioinformatics workflow system based on software containers. The containerisation of software tools provides a wide range of advantages for bioinformatics analyses. Among these benefits, the most important ones are arguably the ability to version and to encapsulate scientific software together with all necessary dependencies within a lightweight portable environment, as required for sound reproducible research. In general it seems clear that the adoption of application containers is gaining momentum in the scientific community. Examples of the adoption of this technology include the BioContainers framework [29] and a number of other projects [30, 31, 12].

As suggested by Burns and Oppenheimer [32], containers are particularly well-suited to being the fundamental "object" in distributed systems. Amongst the variety of available workflow engines, Pachyderm is unique in that it provides an easy mechanism to distribute computations over a collection of containers. Thanks to leveraging multi-node container patterns for distributed algorithms like the scatter/gather pattern, Pachyderm promises what other frameworks such as Apache Spark do [33], but replacing MapReduce-style code [34] or explicit parallelism implementations with legacy code and tools. Moreover, Pachyderm uses upstream Kubernetes as the underlying container scheduler. This use of Kubernetes under-the-hood allows Pachyderm to pass on many of the benefits of Kubernetes to large scale data processing, which have resulted in Kubernetes’ wide spread adoption. These benefits include: (i) optimisation of cluster resource utilisation, (ii) portability between different cloud and on-premise environments, (iii) self-healing of clusters after node/job failures, and (iv) a declarative, unified way of managing applications.

Several workflow tools such as Reflow or Airflow (https://nerds.airbnb.com/airflow) implement specific extensions for each target environment such as Amazon Web Services or Google Cloud. However, Pachyderm leverages Kubernetes for a cloud-native clustering and containerisation orchestration. Thanks to being natively built on Kubernetes, Pachyderm is able to easily distribute and optimally schedule work across nodes using methods such as auto-scaling. On the contrary, Reflow and other workflow tools use their own logic to distribute workloads. Regarding the way workflows are actually specified, Pachyderm pipeline stages are defined in JSON/YAML format that is common within the Kubernetes community, while other workflow tools such as Nextflow implement their own domain specific language (DSL).

Di Tommaso et al. [35] showed that Docker containers have a negligible impact on the performance of a number of bioinformatics pipelines. This study confirms that Pachyderm can scale well despite using application containers. In fact, a scaling efficiency of 79% and a speedup of ∼63 were achieved when using 80 workers in one of the benchmarks. The experiments show that the scaling efficiency decreases as the number of workers is increased. This drop is more significant in the first two benchmarks. An explanation for this can be attributed to a great disparity of running times between the employed input samples. Scientific applications have often changing workload distributions in real life scenarios. These imbalanced workloads among parallel processing units may well result in underutilised computing resources while others are heavily loaded, leading to low overall performance [36].

Despite its numerous advantages, Pachyderm, and in general cloud enabled solutions, have some drawbacks when compared to traditional approaches for biological computations. For instance, despite the fact that our Helm Chart makes it easy to install, Pachyderm is limited to run on Kubernetes, which is not precisely straightforward to set up. Moreover, containerised big data tools such as Pachyderm encounter issues such as (i) limited access to external storage and data locality, (ii) non-optimal container networking and security and (iii) performance overhead when compared to bare-metal settings [37, 38]. Additionally, Pachyderm focuses on data parallelism. This attribute means that, although Pachyderm can process streaming data sources (e.g., from Apache Kafka), it does not offer job parallelism to mitigate the buildup of backpressure in streaming analyses.

Indeed, setting up virtual infrastructures as required for Pachyderm can be quite challenging. In order to show how Pachyderm can be a promising tool for large biological analysis, we created a Helm Chart which simplifies on-demand installations in many types of scenarios, including all popular cloud providers, on-premise and local settings. Moreover, we attempted to demonstrate its suitability by integrating it within the PhenoMeNal Virtual Research Environment and implementing a relatively complex metabolomics workflow and studying its scalability. A major challenge faced when integrating this tool within the VRE was the storage backend, as the PhenoMeNal VRE uses block storage for its services. In contrast, the Pachyderm File System necessitates a cloud-ready object store to interact with the workflow system.

Overall, Pachyderm offers an accessible approach for enabling distributed bioinformatics workflows by using application containers. Likewise, its versioned data management system can be a helpful tool for scientists to keep track of the history of computations and data provenance. All these characteristics, along with being language and cloud agnostic, make it a valuable and powerful tool for creating scalable and reproducible bioinformatics workflows.

